# Metabolic retroconversion of trimethylamine *N*-oxide and the gut microbiota

**DOI:** 10.1101/225581

**Authors:** Lesley Hoyles, Maria L. Jiménez-Pranteda, Julien Chilloux, Francois Brial, Antonis Myridakis, Thomas Aranias, Christophe Magnan, Glenn R. Gibson, Jeremy D. Sanderson, Jeremy K. Nicholson, Dominique Gauguier, Anne L. McCartney, Marc-Emmanuel Dumas

## Abstract

**BACKGROUND:** The dietary methylamines choline, carnitine and phosphatidylcholine are used by the gut microbiota to produce a range of metabolites, including trimethylamine (TMA). However, little is known about the use of trimethylamine *N*-oxide (TMAO) by this consortium of microbes.

**RESULTS:** A feeding study using deuterated TMAO in C57BL6/J mice demonstrated microbial conversion of TMAO to TMA, with uptake of TMA into the bloodstream and its conversion to TMAO. Microbial activity necessary to convert TMAO to TMA was suppressed in antibiotic-treated mice, with deuterated TMAO being taken up directly into the bloodstream. In batch-culture fermentation systems inoculated with human faeces, growth of *Enterobacteriaceae* was stimulated in the presence of TMAO. Human-derived faecal and caecal bacteria (*n* = 66 isolates) were screened on solid and liquid media for their ability to use TMAO, with metabolites in spent media analysed by ^1^H-NMR. As with the *in vitro* fermentation experiments, TMAO stimulated the growth of *Enterobacteriaceae*; these bacteria produced most TMA from TMAO. Caecal/small intestinal isolates of *Escherichia coli* produced more TMA from TMAO than their faecal counterparts. Lactic acid bacteria produced increased amounts of lactate when grown in the presence of TMAO, but did not produce large amounts of TMA. Clostridia (*sensu stricto*), bifidobacteria and coriobacteria were significantly correlated with TMA production in the mixed fermentation system but did not produce notable quantities of TMA from TMAO in pure culture.

**CONCLUSIONS:** Reduction of TMAO by the gut microbiota (predominantly *Enterobacteriaceae*) to TMA followed by host uptake of TMA into the bloodstream from the intestine and its conversion back to TMAO by host hepatic enzymes is an example of metabolic retroconversion. TMAO influences microbial metabolism depending on isolation source and taxon of gut bacterium.

Correlation of metabolomic and abundance data from mixed microbiota fermentation systems did not give a true picture of which members of the gut microbiota were responsible for converting TMAO to TMA; only by supplementing the study with pure culture work and additional metabolomics was it possible to increase our understanding of TMAO bioconversions by the human gut microbiota.

## BACKGROUND

Dietary methylamines such as choline, trimethylamine *N*-oxide (TMAO), phosphatidylcholine (PC) and carnitine are present in a number of foodstuffs, including meat, fish, nuts and eggs. It has long been known that gut bacteria are able to use choline in a fermentation-like process, with trimethylamine (TMA), ethanol, acetate and ATP among the known main end-products [1–3]. TMA can be used by members of the order *Methanomassiliicoccales* (*Archaea*) present in the human gut to produce methane [4], or taken up by the host. Microbially produced TMA derived from PC is absorbed in the small intestine [5]. TMA diffuses into the bloodstream from the intestine via the hepatic vein to hepatocytes, where it is converted to trimethylamine *N*-oxide (TMAO) by hepatic flavin-containing mono-oxygenases (FMOs) [6]. The bulk of TMAO, and lesser amounts of TMA, derived from dietary methylamines can be detected in urine within 6 h of ingestion, and both compounds are excreted in urine but not faeces [7,8]. TMAO can also be detected in human skeletal muscle within 6 h of an oral dose of TMAO [8].

TMAO present in urine and plasma is considered a biomarker for non-alcoholic fatty liver disease (NAFLD), insulin resistance and cardiovascular disease (CVD) [9–12]. Feeding TMAO to high-fat-fed C57BL/6 mice exacerbates impaired glucose tolerance, though the effect on the gut microbiota is unknown [13]. Low plasma PC and high urinary methylamines (including TMAO) were observed in insulin-resistant mice on a high-fat diet, leading to the suggestion microbial methylamine metabolism directly influences choline bioavailability; however, the microbiota of the mice was not studied nor was labeled choline used to determine the metabolic fate of TMA/TMAO [9]. Choline bioavailability is thought to contribute to hepatic steatosis in patients [11]. Spencer *et al.* [11] manipulated dietary choline levels in 15 females for 2 weeks and monitored changes in the fecal microbiota during the intervention. They found the level of choline supplementation affected Gammaproteobacteria (inhibited by high levels of dietary choline, negatively correlated with changes in liver fat/spleen fat ratios) and Erysipelotrichi (positively correlated with changes in liver fat/spleen fat ratios), suggesting baseline levels of these taxa may predict the susceptibility of an individual to liver disease. However, the study was under-powered and requires replication with a much larger cohort to fully examine the link between choline bioavailability and liver disease. High levels of circulating TMAO are associated with CVD [12], potentially via a platelet-mediated mechanism [14]. Circulating TMAO has been suggested to play a role in protection from hyperammonemia in mice, acting as an osmoprotectant, and from glutamate neurotoxicity in mice and primary cultures of neurons [15,16]. Recently, chronic exposure to TMAO was shown to attenuate diet-associated impaired glucose tolerance in mice, and reduce endoplasmic reticulum stress and adipogenesis in adipocytes [10]. The relevance of these findings to human health remains to be determined. There is a basal level of TMA and TMAO detected in human urine even in the absence of dietary supplementation [17], suggesting use of (microbial and/or host) cell-derived choline or PC in the intestinal tract by the gut microbiota.

While it is well established that choline [2,11,18], PC [2,12] and carnitine [19–22] are used by the human and rodent gut microbiotas to produce TMA, little is known about the reduction of TMAO (predominantly from fish) to TMA (or other compounds) by members of these consortia. TMAO is found at concentrations of 20–120 mg per 100 g fish fillet [23]. Ingestion of fish by humans leads to increased urinary excretion of dimethylamine (DMA) and TMA, from 5.6 to 24.1 and from 0.2 to 1.6 μmol/24 h/kg of body weight, respectively [24]. Individuals with trimethylaminuria (fish odour syndrome), in which TMA is not detoxified to TMAO by hepatic FMOs, have been shown to reduce 40–60 % of an oral dose of TMAO to TMA [25]. It was suggested the gut microbiota was responsible for reducing TMAO in these individuals, and in those individuals without trimethylaminuria the TMA was re-oxidized in the liver before being excreted in the urine. The process was termed ‘*metabolic retroversion*’ “*(i.e., a cycle of reductive followed by oxidative reactions to regenerate TMAO)*” [25]. The reduction of TMAO to TMA is most commonly associated with the gut microbiota of marine fish, with the TMA generated by these bacteria contributing to the characteristic odour of rotting fish [26]. Members of the class *Gammaproteobacteria*, which includes a number of food spoilage organisms, are known to reduce TMAO to TMA quantitatively and are unable to reduce TMA further [26,27]. The conversion of TMAO to TMA by bacteria represents a unique form of anaerobic respiration, in which TMAO reductase acts as a terminal electron acceptor (from NADH or formate) for respiratory electron flow [27].

Mining of metagenomic data (*torA*) suggests *Proteobacteria* (particularly *Escherichia* and *Klebsiella* spp.) are likely to contribute greatest to the production of TMA from TMAO in the human gut via the TMAO reductase pathway, with *Actinobacteria* (*Eggerthellaceae*) becoming more important under stress [28]. Other microbial genes are associated with production of TMA from choline (*cutC*), glycine-betaine (*grdH*), L-carnitine (*cntA*) and γ-butyrobetaine (*cntA*) [4]. A search of the NCBI nucleotide database with the phrase ‘TMAO reductase’ also suggests many other bacteria of human intestinal origin (namely, *Salmonella, Helicobacter, Prevotella, Bacillus* and *Bacteroides* spp.) should be able to reduce TMAO to TMA. However, this trait has not been examined *in vitro* for isolates of intestinal origin.

Consequently, we carried out an *in vivo* study in mice to confirm use of TMAO by the mouse gut microbiota and to allow us to examine metabolic retroconversion of TMAO in a murine model. We then used an *in vitro* fermentation system to highlight the effect of TMAO on the human faecal microbiota. Finally, we screened a panel of human faecal and caecal/small intestinal isolates to determine which members of the human gut microbiota were able to reduce TMAO to TMA, and whether their metabolism was affected by being grown in the presence of TMAO.

## METHODS

### Animal work

Groups of six-week-old C57BL6/J mice (Janvier Labs, Courtaboeuf, France) were received and acclimated in specific pathogen-free (SPF) maintenance conditions. They were fed with a standard chow diet (R04-40, Safe, Augy, France) and were either given free access to tap water or tap water containing the antibiotic cocktail (0.5 g/L vancomycin hydrochloride, 1 g/L neomycin trisulfate, 1 g/L metronidazole, 1 g/L ampicillin sodium; all antibiotics were purchased from Sigma-Aldrich) for 14 days. Mice were given in the morning by gavage either a solution of d_9_-TMAO at 1×10^-4^ M (Cambridge Isotope Laboratories Inc., DLM-4779-0, UK) or saline and euthanized 6 h later by cervical dislocation. Blood samples were collected by tail tipping every 2 h in Microvette^®^ CB 300 Lithium Heparin (Sarstedt, Marnay, France). Plasma was separated by centrifugation (10 min, 5000 ***g***, 4 °C) and stored at −80 °C until analysed by ultra-performance liquid chromatography–tandem mass spectrometry (UPLC–MS/MS).

### UPLC–MS/MS determination of plasma TMA, d_9_-TMA, TMAO and d_9_-TMAO

UPLC–MS/MS was employed for the determination of TMA, d_9_-TMA, TMAO and d_9_-TMAO. Samples (10 µL) were spiked with 10 µL Internal Standard solution (d_9_-choline and ^13^C_3_/^15^N-TMA in water; 1 mg/L, Sigma-Aldrich). Ethyl 2-bromoacetate solution (45 µL) (15 g/L ethyl 2-bromoacetate, 1 % NH_4_OH in acetonitrile; ChromaSolv grade, Sigma-Aldrich) was added and derivatization of TMAs (TMA, d_9_-TMA and ^13^C_3_/^15^N-TMA) to their ethoxy-analogues was completed after 30 min at room temperature. Protein/lipid precipitation solution (935 µL) (94 % acetonitrile/5 %water/1 % formic acid; ChromaSolv grade, Sigma-Aldrich) was added, samples were centrifuged (20 min, 20,000 ***g***, 4 °C) and were transferred to UPLC-autosampler vials. Sample injections (5 µL loop) were performed with a Waters Acquity UPLC-Xevo TQ-S UPLC–MS/MS system equipped with an Acquity BEH HILIC (2.1×100 mm, 1.7 µm) chromatographic column. An isocratic elution was applied with 10 mM ammonium formate (Sigma-Aldrich) in 95:5 (v/v) acetronitrile:water for 7 min at 750 µL/min and 50 °C. Positive electrospray (ESI^+^) was used as ionization source and mass spectrometer parameters were set as follows: capillary, cone and sources voltages at −700, −18 and 50 V, respectively, desolvation temperature at 600 °C, desolvation/cone/nebuliser gases were high purity nitrogen (BOC) at 1000 L/h, 150 L/h and 7 bar, respectively. Collision gas was high-purity argon (BOC). Mass spectrometer was operated in multiple reaction monitoring mode. The monitored transitions were the following: for TMA, +146 → +118/59 *m*/*z* (23/27 V); for d_9_-TMA, +155 → +127/68 *m*/*z* (21/23 V); for ^13^C_3_/^15^N-TMA, +150 → +63/122 *m*/*z* (27/22 V); for TMAO, +76 → +59/58 *m*/*z* (12/13 V); for d_9_-TMAO, +85 → +68/66 *m*/*z* (18/20 V); and for d_9_-choline, +108 → +60/45 *m*/*z* (20/22 V). The system was controlled by MassLynx software, also used for data acquisition and analysis.

### ^1^H-NMR spectroscopy and data analysis

Medium samples were randomized and centrifuged (5 min, 16000 ***g***). Aliquots (50 µL) of the supernatants were diluted in 550 µL D_2_O containing 1 mM trimethylsilyl-(2,2,3,3-2H_4_)-1-propionate. Samples were transferred to 5 mm NMR tubes and measured on a NMR spectrometer (Bruker) operating at 600.22 MHz ^1^H frequency as described [29]. ^1^H-NMR spectra were pre-processed and analysed as described [9] using the Statistical Recoupling of Variables-algorithm [30]. Structural assignment was performed as described [31], using in-house and publicly available databases.

### *In vitro* fermentation systems

Freshly voided faecal samples were obtained from three healthy human volunteers (one male and two females; age range 29–31 years), none of whom had been prescribed antibiotics 6 months prior to the study, eaten fish 3 days prior to sample collection or had any history of gastrointestinal disease. The University of Reading’s Research Ethics Committee (UREC) does not require that specific ethical review and approval be given by UREC for the collection of faecal samples from healthy human volunteers to inoculate *in vitro* fermentation systems. Samples were processed immediately by diluting them 1 in 10 (w/w) in pre-reduced minimal broth and homogenizing them in a filter bag in a stomacher (Stomacher 400 Lab System; Seward) for 2 min at ‘high’ speed. For each donor, a batch culture vessel containing 175 mL of pre-reduced minimal broth was inoculated with 20 mL of the faecal homogenate and 5 mL of sterile H_2_O containing 2 g TMAO dihydrate (Sigma-Aldrich). Control vessels were inoculated with 20 mL of the faecal homogenate and 5 mL of sterile H_2_O. The final working volume of each batch culture vessel was 200 mL. The pH of each vessel (pH 6.5) was controlled automatically by the addition of 2 M HCl or 2 M NaOH. pH controllers were supplied by Electrolab. The contents of each vessel were stirred constantly. An anaerobic environment was maintained by constantly sparging the vessels with O_2_-free N_2_. The temperature of each vessel was maintained at 37 °C by use of a circulating waterbath connected to each fermentation vessel. The experiment was run for 9 h, with samples taken at 0, 1, 2, 3, 4, 5, 6 and 9 h.

### Fluorescence *in situ* hybridization (FISH) analysis

Aliquots (2× 375 μL) of sample were fixed in ice-cold 4 % paraformaldehyde for 4 h, washed in sterile phosphate-buffered saline and stored for FISH analysis as described [32]. **Supplementary Table 1** gives details for probes used in this study. Probes were synthesized commercially (MWG-Biotech) and labelled with the fluorescent dye cyanine 3 (Cy3; excitation λ, 514 nm; emission λ, 566 nm; fluorescence colour, orange–red). FISH was done as described [32].

Slides were viewed under a Nikon E400 Eclipse microscope. DAPI slides were visualized with the aid of a DM400 filter; probe slides were visualized with the aid of a DM575 filter. Cells (between 15 and 50 per field of view) were counted for 15 fields of view, and the numbers of bacteria were determined by using the following equation:

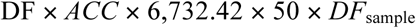

Where the DF (dilution factor) was calculated by taking into account the concentration of the original sample (375 μL to 300 μL = 0.8×). *ACC* (average cell count) was determined by counting 15 fields of view and assumes that a normal distribution was observed for the counts. The figure 6,732.42 refers to the area of the well divided by the area of the field of view. *DF*_sample_ refers to the dilution of sample used with a particular probe (e.g. 5× for *t*_0_ Lab158 counts). The detection limit of this method was 89,766 bacteria/mL of sample (= log_10_ 4.95).

### Screening of bacteria for ability to reduce TMAO

Part of an in-house collection (University of Reading) of bacteria isolated from human caecal/small intestinal and faecal samples (**Table 1**) was screened on minimal agar [g/L: glucose (Fisher Scientific), 4; bacteriological peptone (Oxoid), 20; NaCl (Fisher Scientific), 5; neutral red (Sigma-Aldrich), 0.03; agar technical no. 3 (Oxoid), 15; pH 7] with and without 1 % (w/v) TMAO dihydrate (Sigma-Aldrich). The composition of the agar was based on [27], who used the medium with crystal violet and bile salts to screen members of the *Enterobacteriaceae* for TMAO reductase activity. Colonies of isolates able to ferment glucose were red, whereas those able to reduce TMAO were white. Growth curves (OD_600_ measured hourly for 10–12 h, then at 24 h) were determined for selected isolates grown anaerobically at 37 °C in anaerobic minimal broth [g/L: glucose, 4; bacteriological peptone, 20; NaCl, 5; L-cysteine HCl (Sigma-Aldrich), 0.5; resazurin solution (0.25 mg/mL; Sigma-Aldrich), 4 mL; pH 7] with and without 1 % (w/v) TMAO dihydrate. Glucose was substituted by raffinose (Sigma-Aldrich) [33] in the minimal broth when working with bifidobacteria because of the poor growth of these bacteria on the glucose-based medium. pH and metabolite profiles of culture medium were examined at the end of the growth experiment (i.e. at *t*_24_).

**Table 1.**
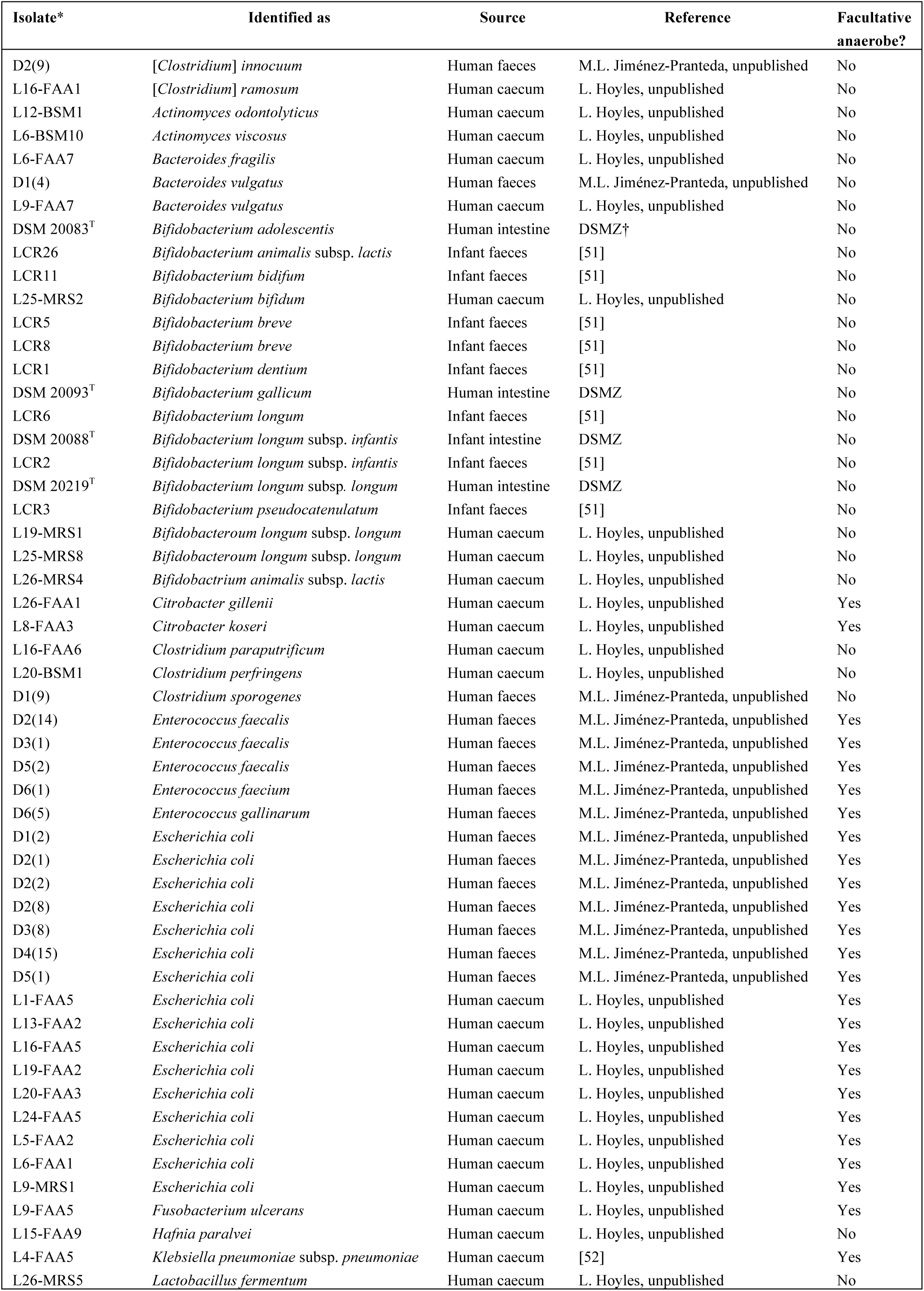
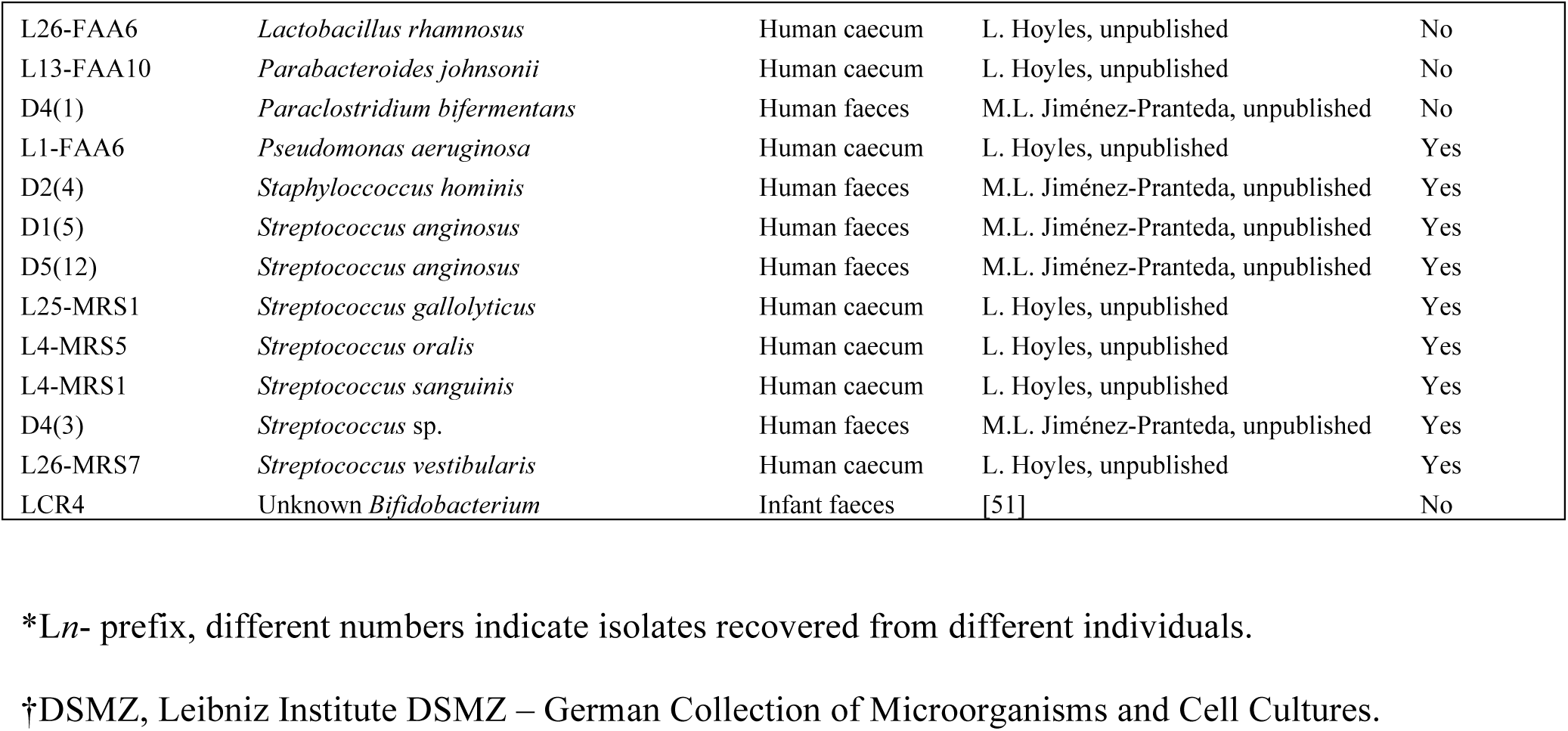
Details for human-derived gut bacteria screened for their ability to reduce or utilize TMAO.

| Isolate* | Identified as | Source | Reference | Facultative anaerobe? |
| --- | --- | --- | --- | --- |
| D2(9) | *[Clostridium] innocuum* | Human faeces | M.L. Jiménez-Pranteda, unpublished | No |
| L16-FAA1 | *[Clostridium] ramosum* | Human caecum | L. Hoyles, unpublished | No |
| L12-BSM1 | *Actinomyces odontolyticus* | Human caecum | L. Hoyles, unpublished | No |
| L6-BSM10 | *Actinomyces viscosus* | Human caecum | L. Hoyles, unpublished | No |
| L6-FAA7 | *Bacteroides fragilis* | Human caecum | L. Hoyles, unpublished | No |
| D1(4) | *Bacteroides vulgatus* | Human faeces | M.L. Jiménez-Pranteda, unpublished | No |
| L9-FAA7 | *Bacteroides vulgatus* | Human caecum | L. Hoyles, unpublished | No |
| DSM 20083<sup>T</sup> | *Bifidobacterium adolescentis* | Human intestine | DSMZ<sup>†</sup> | No |
| LCR26 | *Bifidobacterium animalis* subsp. *lactis* | Infant faeces | [51] | No |
| LCR11 | *Bifidobacterium bidifum* | Infant faeces | [51] | No |
| L25-MRS2 | *Bifidobacterium bifidum* | Human caecum | L. Hoyles, unpublished | No |
| LCR5 | *Bifidobacterium breve* | Infant faeces | [51] | No |
| LCR8 | *Bifidobacterium breve* | Infant faeces | [51] | No |
| LCR1 | *Bifidobacterium dentium* | Infant faeces | [51] | No |
| DSM 20093<sup>T</sup> | *Bifidobacterium gallicum* | Human intestine | DSMZ | No |
| LCR6 | *Bifidobacterium longum* | Infant faeces | [51] | No |
| DSM 20088<sup>T</sup> | *Bifidobacterium longum* subsp. *infantis* | Infant intestine | DSMZ | No |
| LCR2 | *Bifidobacterium longum* subsp. *infantis* | Infant faeces | [51] | No |
| DSM 20219<sup>T</sup> | *Bifidobacterium longum* subsp. *longum* | Human intestine | DSMZ | No |
| LCR3 | *Bifidobacterium pseudocatenulatum* | Infant faeces | [51] | No |
| L19-MRS1 | *Bifidobacterium longum* subsp. *longum* | Human caecum | L. Hoyles, unpublished | No |
| L25-MRS8 | *Bifidobacterium longum* subsp. *longum* | Human caecum | L. Hoyles, unpublished | No |
| L26-MRS4 | *Bifidobacterium animalis* subsp. *lactis* | Human caecum | L. Hoyles, unpublished | No |
| L26-FAA1 | *Citrobacter gillenii* | Human caecum | L. Hoyles, unpublished | Yes |
| L8-FAA3 | *Citrobacter koseri* | Human caecum | L. Hoyles, unpublished | Yes |
| L16-FAA6 | *Clostridium paraputrificum* | Human caecum | L. Hoyles, unpublished | No |
| L20-BSM1 | *Clostridium perfringens* | Human caecum | L. Hoyles, unpublished | No |
| D1(9) | *Clostridium sporogenes* | Human faeces | M.L. Jiménez-Pranteda, unpublished | No |
| D2(14) | *Enterococcus faecalis* | Human faeces | M.L. Jiménez-Pranteda, unpublished | Yes |
| D3(1) | *Enterococcus faecalis* | Human faeces | M.L. Jiménez-Pranteda, unpublished | Yes |
| D5(2) | *Enterococcus faecalis* | Human faeces | M.L. Jiménez-Pranteda, unpublished | Yes |
| D6(1) | *Enterococcus faecium* | Human faeces | M.L. Jiménez-Pranteda, unpublished | Yes |
| D6(5) | *Enterococcus gallinarum* | Human faeces | M.L. Jiménez-Pranteda, unpublished | Yes |
| D1(2) | *Escherichia coli* | Human faeces | M.L. Jiménez-Pranteda, unpublished | Yes |
| D2(1) | *Escherichia coli* | Human faeces | M.L. Jiménez-Pranteda, unpublished | Yes |
| D2(2) | *Escherichia coli* | Human faeces | M.L. Jiménez-Pranteda, unpublished | Yes |
| D2(8) | *Escherichia coli* | Human faeces | M.L. Jiménez-Pranteda, unpublished | Yes |
| D3(8) | *Escherichia coli* | Human faeces | M.L. Jiménez-Pranteda, unpublished | Yes |
| D4(15) | *Escherichia coli* | Human faeces | M.L. Jiménez-Pranteda, unpublished | Yes |
| D5(1) | *Escherichia coli* | Human faeces | M.L. Jiménez-Pranteda, unpublished | Yes |
| L1-FAA5 | *Escherichia coli* | Human caecum | L. Hoyles, unpublished | Yes |
| L13-FAA2 | *Escherichia coli* | Human caecum | L. Hoyles, unpublished | Yes |
| L16-FAA5 | *Escherichia coli* | Human caecum | L. Hoyles, unpublished | Yes |
| L19-FAA2 | *Escherichia coli* | Human caecum | L. Hoyles, unpublished | Yes |
| L20-FAA3 | *Escherichia coli* | Human caecum | L. Hoyles, unpublished | Yes |
| L24-FAA5 | *Escherichia coli* | Human caecum | L. Hoyles, unpublished | Yes |
| L5-FAA2 | *Escherichia coli* | Human caecum | L. Hoyles, unpublished | Yes |
| L6-FAA1 | *Escherichia coli* | Human caecum | L. Hoyles, unpublished | Yes |
| L9-MRS1 | *Escherichia coli* | Human caecum | L. Hoyles, unpublished | Yes |
| L9-FAA5 | *Fusobacterium ulcerans* | Human caecum | L. Hoyles, unpublished | Yes |
| L15-FAA9 | *Hafnia paralvei* | Human caecum | L. Hoyles, unpublished | No |
| L4-FAA5 | *Klebsiella pneumoniae* subsp. *pneumoniae* | Human caecum | [52] | Yes |
| L26-MRS5 | *Lactobacillus fermentum* | Human caecum | L. Hoyles, unpublished | No |
| L26-FAA6 | *Lactobacillus rhamnosus* | Human caecum | L. Hoyles, unpublished | No |
| L13-FAA10 | *Parabacteroides johnsonii* | Human caecum | L. Hoyles, unpublished | No |
| D4(1) | *Paraclostridium bifermentans* | Human faeces | M.L. Jiménez-Pranteda, unpublished | No |
| L1-FAA6 | *Pseudomonas aeruginosa* | Human caecum | L. Hoyles, unpublished | Yes |
| D2(4) | *Staphylococcus hominis* | Human faeces | M.L. Jiménez-Pranteda, unpublished | Yes |
| D1(5) | *Streptococcus anginosus* | Human faeces | M.L. Jiménez-Pranteda, unpublished | Yes |
| D5(12) | *Streptococcus anginosus* | Human faeces | M.L. Jiménez-Pranteda, unpublished | Yes |
| L25-MRS1 | *Streptococcus gallolyticus* | Human caecum | L. Hoyles, unpublished | Yes |
| L4-MRS5 | *Streptococcus oralis* | Human caecum | L. Hoyles, unpublished | Yes |
| L4-MRS1 | *Streptococcus sanguinis* | Human caecum | L. Hoyles, unpublished | Yes |
| D4(3) | *Streptococcus* sp. | Human faeces | M.L. Jiménez-Pranteda, unpublished | Yes |
| L26-MRS7 | *Streptococcus vestibularis* | Human caecum | L. Hoyles, unpublished | Yes |
| LCR4 | Unknown *Bifidobacterium* | Infant faeces | [51] | No |

\*Ln- prefix, different numbers indicate isolates recovered from different individuals.
†DSMZ, Leibniz Institute DSMZ – German Collection of Microorganisms and Cell Cultures.

### Statistical analyses

Differences between metabolites produced by faecal and caecal/small intestinal isolates of *Escherichia coli* and lactic acid bacteria in the presence and absence of TMAO were analysed using Student’s *t* test. FISH data from the *in vitro* fermentation systems were analysed using the Kolmogorov–Smirnov test with statistical significance, after correction for multiple testing, taken at *P* < 0.05. Because of the presence of ties in the metabolite data from *in vitro* fermentation systems, the bootstrapped Kolmogorov–Smirnov test (10,000 replications) was used with these data. Data from *in vitro* fermentation systems (FISH and NMR) were correlated using Spearman’s rank correlation (corrected for ties) with results corrected for multiple testing using the method of Benjamini and Hochberg [34].

## RESULTS

### *In vivo* confirmation of metabolic retroconversion

First, to confirm *in vivo* metabolic retroconversion (i.e. microbial conversion of TMAO to TMA, followed by host conversion of TMA to TMAO), we administered isotopically-labelled d_9_-TMAO or saline to mice that had or had not been treated with a broad-spectrum antibiotic cocktail for 14 days to suppress the gut microbiota [12]. Reduction of d_9_-TMAO to d_9_-TMA was quantified in urine by UPLC–MS/MS up to 6 h after d_9_-TMAO gavage, together with unlabelled TMA and TMAO potentially produced from dietary sources (**Fig. 1**).

**Fig. 1.**
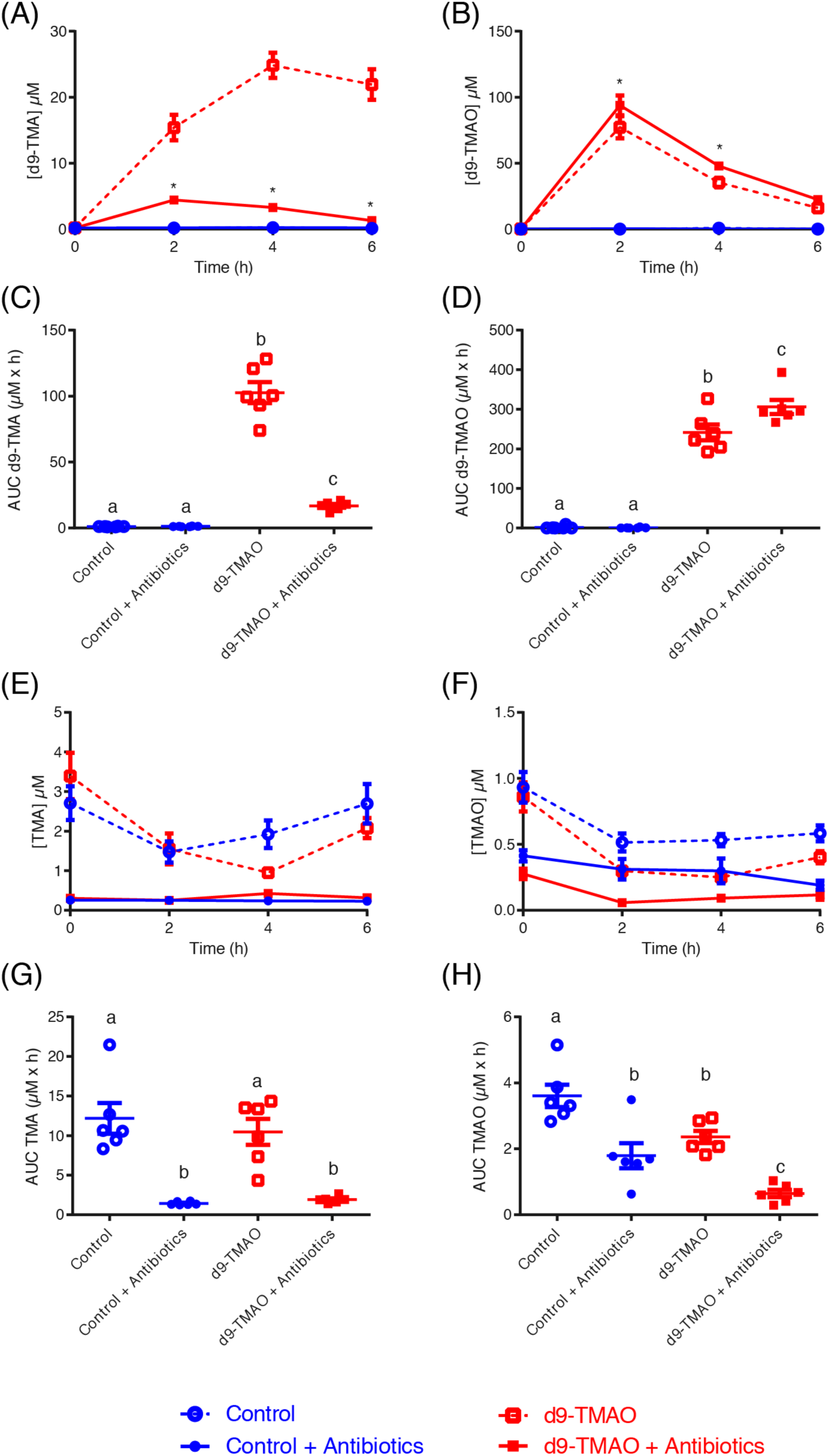
*In vivo* confirmation of metabolic retroconversion of TMAO. Reduction of d_9_-TMAO to d_9_-TMA was quantified by UPLC–MS/MS up to 6 h after d_9_-TMAO gavage and antibiotic treatment, together with unlabelled TMA and TMAO levels. Plasma quantification of post-gavage (A) d_9_-TMA and (B) d_9_-TMAO. *, Significantly (*P* < 0.05; *t* test and corrected for multiple comparison using the Holm–Sidak method) different from the respective groups not treated with antibiotics. (C) d_9_-TMA bioavailability (AUC). (D) d_9_-TMAO bioavailability (AUC). Plasma quantification of post-gavage unlabelled/endogenous (E) TMA and (F) TMAO. *, Significant between d_9_ and d_9_ antibiotic treatment; $, significant between TMAO and TMAO antibiotic treatment. (G) Unlabelled/endogenous TMA bioavailability (AUC). (H) Unlabelled/endogenous TMAO bioavailability (AUC). Data (*n* = 6 per group) are shown as mean ± SEM. (A, B, E and F). Differences between the bioavailabilities (C, D, G and H) were assessed using one-way analysis of variance (ANOVA), followed by Holm–Sidak post hoc tests. Data with different superscript letters are significantly different (*P* < 0.05).

In the absence of antibiotics, d_9_-TMAO was converted to d_9_-TMA within 2 h of gavage. This conversion was dramatically reduced, leading to a significantly lower concentration of d_9_-TMA and a higher concentration of d_9_-TMAO, when the gut microbiota was suppressed by antibiotics (**Fig. 1A–D**). We noted that in the absence of antibiotics all animals excreted approximately three times higher levels of urinary unlabelled TMA than TMAO (**Fig. 1E–H**), corresponding to constitutively low FMO3 activity in mice [35]. Levels of unlabelled TMAO/TMA were not significantly different from one another in the control and d_9_-TMAO-fed animals, while antibiotic treatment significantly reduced the amount of both TMAO and TMA (**Fig. 1E, F**). Bioavailability of unlabelled TMAO/TMA, as assessed by area under the curve (AUC), was significantly reduced by antibiotic treatment in both saline and d_9_-TMAO-gavaged animals, with almost no TMA detected in either experimental group (**Fig. 1G, H**).

### Effect of TMAO on human gut bacteria within a mixed system

After *in vivo* validation of the role of the gut microbiota in metabolic retroconversion, we analysed the effect of TMAO on the faecal microbiota in an anaerobic batch-culture fermentation system. Fermenter vessels filled with the glucose-containing medium supplemented or not with 1 % (w/v) TMAO were inoculated with faecal slurries from three healthy donors and monitored for 9 h. With the exception of enhanced growth of the *Enterobacteriaceae* (probe Ent), the presence of TMAO in the medium had no statistically significant effect on the growth of bacteria within the fermentation systems at 9 h (**Fig. 2A, Supplementary Figure 1**).

**Fig. 2.**
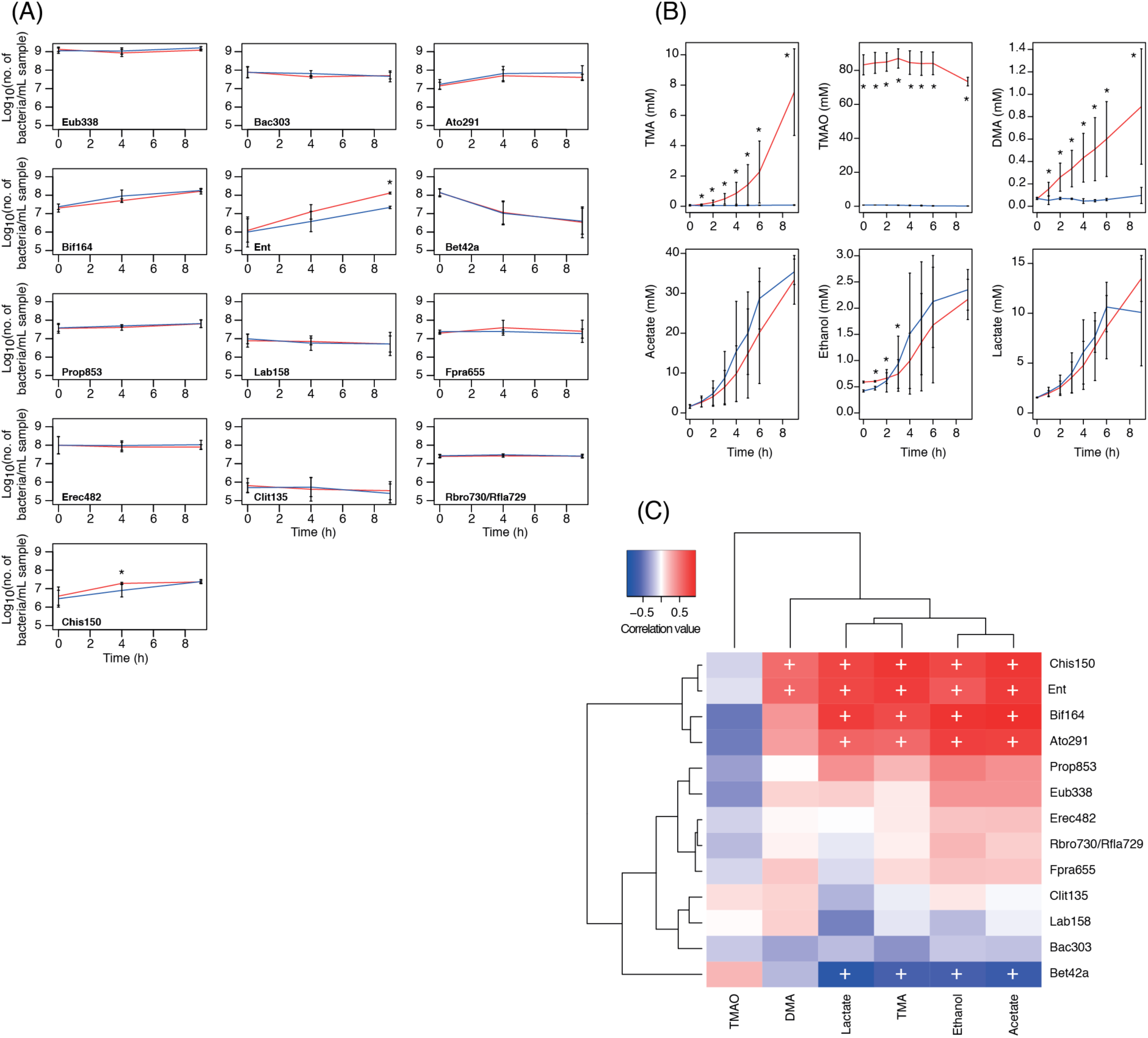
Effect of TMAO on mixed faecal microbial population *in vitro*. (A) Enumeration of selected bacteria in fermentation vessels by FISH analysis. Red lines, TMAO-containing systems; blue lines, negative controls. Data are shown as mean + SD (*n* = 3). Eub338, total bacteria; Ent, *Enterobacteriaceae*; Bif164, *Bifidobacterium* spp.; Lab158, lactic acid bacteria. *, Statistically significantly different (adjusted *P* < 0.05) from the control at the same time point. Full data are shown in **Supplementary Figure 1**. (B) ^1^H-NMR data for batch culture samples. Data are shown mean ± SD (*n* = 3). Red lines, TMAO-containing systems; blue lines, negative controls. *, Statistically significantly different (*P* < 0.05) from the negative control at the same time point. (C) Bidirectional clustering of correlation matrix of FISH data and data for the six metabolites found in highest amounts in the NMR spectra from the batch-culture samples. +, Adjusted *P* value (Benjamini– Hochberg) statistically significant (*P* < 0.05). FISH and metabolite data and a table of correlations and adjusted *P* values (Benjamini–Hochberg) for the batch-culture samples are available in **Supplementary Tables 3–5**.

Huge variability, as measured by ^1^H-NMR, was observed in the amount of TMA and DMA produced by gut bacteria in the TMAO-containing fermentation systems, with the concentrations of both metabolites increasing steadily from 0 to 9 h and differing significantly (*P* < 0.05) from the control systems (**Fig. 2B**). The amount of TMAO in the systems was seen to decrease at 8 h.

Correlation of metabolite and FISH data demonstrated *Clostridium* clusters I and II (Chis150), *Enterobacteriaceae* (Ent), bifidobacteria (Bif164) and coriobacteriia (Ato291) were associated with TMA, acetate, ethanol and lactate (**Fig. 2C**). The *Betaproteobacteria* (Bet42a) were anti-correlated with TMA, acetate, ethanol and lactate, which is unsurprising given this was the only group of bacteria whose representation decreased in the fermentation systems over the course of the experiment (**Supplementary Figure 1, Fig. 2C**). The *Enterobacteriaceae* and clostridia were positively correlated with DMA production. The lactic acid bacteria (Lab158) were not significantly correlated with any of the metabolites in the mixed culture.

### Growth of pure cultures of gut bacteria in the presence of TMAO

Using an in-house collection of bacteria, we initially screened isolates on a modified version of the agar of Takahi & Ishimoto [27] as a rapid means of screening bacteria for TMAO reductase activity, and thereby their ability to reduce TMAO to TMA. While members of the *Enterobacteriaceae* produced expected results (i.e. white colonies when grown in the presence of TMAO and glucose, rather than red colonies on the glucose only control), we observed a number of unexpected outcomes depending on the species under study. For example, colonies of lactic acid bacteria were larger (almost twice their usual size) on TMAO-containing agar than on the glucose control but remained red in colour, suggesting they had not reduced TMAO to TMA at detectable levels but TMAO was influencing their growth. The clostridia examined produced mixed results on the control and TMAO-containing media (i.e. white colonies on both plates or on the control plate only, larger colonies on the TMAO-containing medium but without a colour change of the medium).

To determine whether these isolates were converting TMAO to TMA but a low level, we examined the growth of all isolates in liquid culture and metabolites in the spent medium using NMR. The growth of the *Enterobacteriaceae* was most greatly affected by the presence of TMAO in the medium, with a faster, longer-lasting exponential phase than for the same isolates grown in the control medium (**Fig. 3A**). pH of the spent medium (after 24 h) when *Enterobacteriaceae* were grown in the presence of TMAO increased from a mean of 4.7 ± 0.3 (for the control) to 7.6 ± 0.3 (*n* = 20) (the change in pH is what causes the colonies to appear white on TMAO-containing agar). The growth of lactic acid bacteria, including *Enterococcus* and *Streptococcus* (**Fig. 3A**) spp., was enhanced in the presence of TMAO, but not to the same extent as seen for the *Enterobacteriaceae*. There was no significant difference (*P* = 0.27, *t* test) in the pH of the spent medium for these bacteria after 24 h (mean 4.67 ± 0.9 compared with 4.33 ± 0.27 for the control). The growth of members of *Clostridium* cluster I (e.g. *Clostridium perfringens*, **Fig. 3A**) was not enhanced in the presence of TMAO, though some of these bacteria changed the colour of both the control and TMAO-containing plates yellow during their growth. The pH of the spent liquid medium confirmed this observation to be due to the alkalinity of the media in the control and TMAO-containing media after 24 h incubation: e.g. *Clostridium sporogenes* D1(9) (pH 6.25 compared with 6.76 in the control medium), *Clostridium paraputrificum* L16-FAA6 (pH 6.45 vs 5.38) and *Clostridium perfringens* L20-BSM1 (pH 5.56 vs 4.64).

**Fig. 3.**
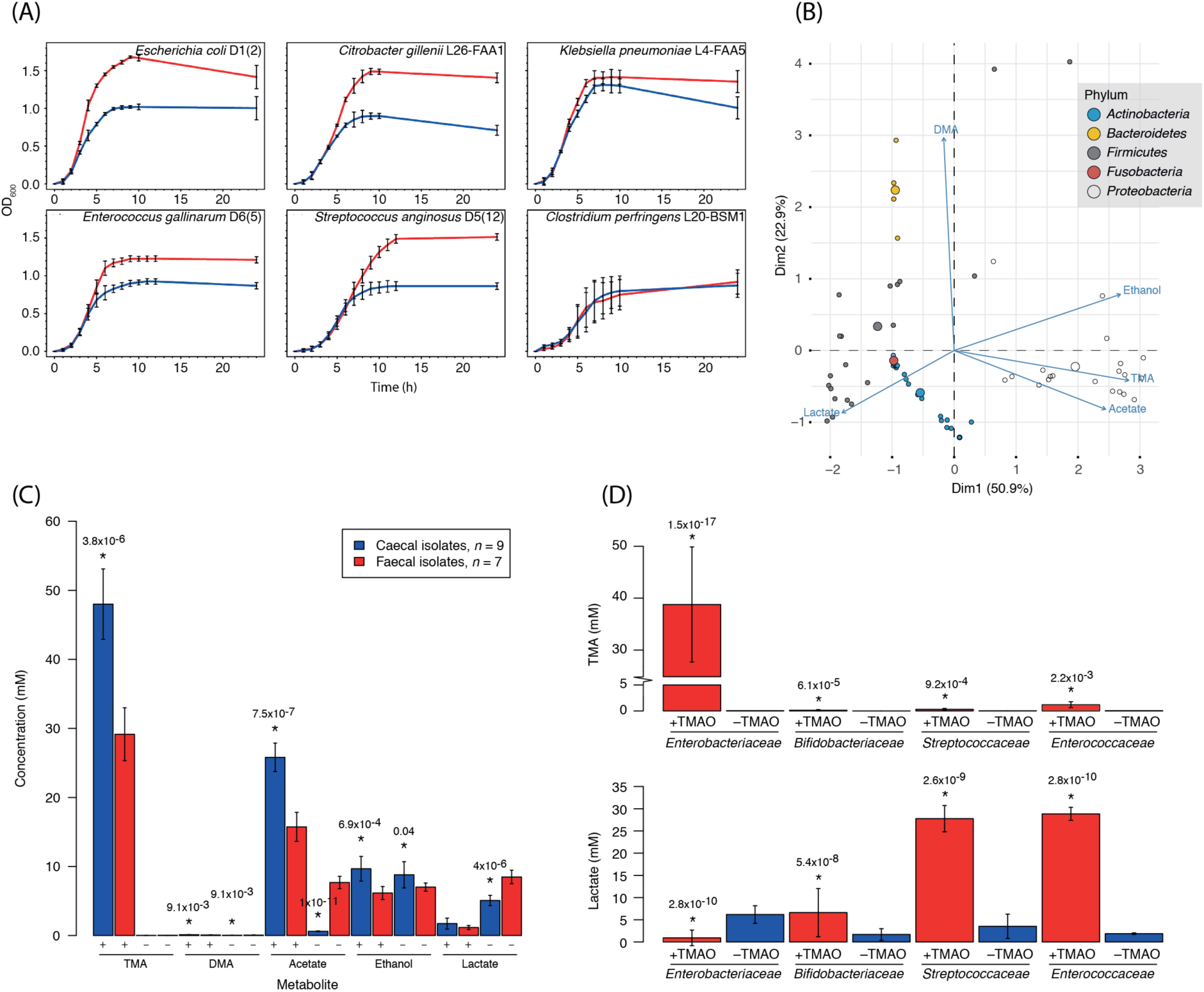
Influence of TMAO on growth and metabolism of pure cultures of gut bacteria. (A) Representative growth curves for isolates grown in the presence and absence of TMAO. Red lines, TMAO-supplemented cultures; blue lines, negative controls. Data are shown as mean ± SD (*n* = 3). (B) Biplot showing production of various metabolites when isolates were grown in the presence of TMAO. Summary of data from **Supplementary Table 2**. The larger a circle, the more of the metabolite produced by an isolate. (C) Differences in metabolites produced when caecal and faecal isolates of *Escherichia coli* were grown in the presence (+) and absence (–) of 1 % TMAO. Adjusted (Benjamini–Hochberg) *P* values indicate the caecal isolates were significantly different from the faecal isolates for a particular metabolite. (D) Lactate production by lactic acid bacteria was increased in the presence of TMAO. *Enterobacteriaceae, n* = 20; *Bifidobacteriaceae, n* = 17; *Streptococcaceae, n* = 7; *Enterococcaceae, n* = 5. Members of the *Enterococcaceae* and *Streptococcaceae* are homofermenters (produce only lactic acid from glucose fermentation), whereas the *Bifidobacteriaceae* are heterofermenters (produce ethanol, CO_2_ and lactic acid from glucose fermentation), though it should be noted the bifidobacteria included in this study were grown on raffinose-containing media. Red, TMAO-containing medium; blue, negative control. *, Statistically significantly different from its negative control (adjusted *P* value < 0.05).

^1^H-NMR analysis of spent medium from the TMAO-containing and control samples demonstrated that, as expected, the *Enterobacteriaceae* produced the greatest amount of TMA from TMAO (mean 38.79 ± 11.08 mM compared with 0.03 ± 0.01 mM, *n* = 20) (**Fig. 3B**; **Supplementary Table 2**). Members of the families *Peptostreptococcaceae* (3.72 mM, *n* = 1), *Clostridiaceae* (Cluster I) (2.62 ± 1.83 mM, *n* = 3), *Porphyromonadaceae* (1.42 mM, *n* = 1), *Bacteroidaceae* (1.40 ± 0.31 mM, *n* = 3), *Enterococcaceae* (1.19 ± 0.05 mM, *n* = 5), *Erysipelotrichaceae* (0.94 mM, *n* = 1; [*Clostridium*] *ramosum*), *Staphylococcaceae* (0.34 mM, *n* = 1), *Streptococcaceae* (0.30 ± 0.16 mM, *n* = 5), *Lactobacillaceae* (0.17 ± 0.07 mM, *n* = 2), *Pseudomonadaceae* (0.12 mM, *n* = 1) and *Bifidobacteriaceae* (0.13 ± 0.1 mM, *n* = 17) produced low levels of TMA from TMAO (**Fig. 3B**; **Supplementary Table 2**). There was great variability in the ability of the bifidobacteria to produce TMA from TMAO, with several isolates and [*Clostridium*] *innocuum, Actinomyces odontolyticus, Fusobacterium ulcerans* and *Actinomyces viscosus* not producing TMA from TMAO (**Fig. 3B**; **Supplementary Table 2**).

### Differences in the metabolic capabilities of faecal and caecal *Escherichia coli* isolates

Comparison of the amounts of TMA, and co-metabolites, produced by the faecal (*n* = 7) and caecal (*n* = 9) isolates of *Escherichia coli* demonstrated significantly higher amounts of TMA were produced by the caecal isolates compared with the faecal isolates in the TMAO-containing medium (**Fig. 3C**). *Escherichia coli* of caecal origin produced more TMA than faecal isolates of the same bacterium or other enterobacteria (*Hafnia, Citrobacter* and *Klebsiella* spp.) (**Supplementary Table 2**). The faecal isolates produced more acetate and lactate than the caecal isolates when grown in the control medium. Taken together, these results demonstrate the different metabolic capabilities of isolates of *Escherichia coli* recovered from different regions of the human gut.

### Lactic acid bacteria produce more lactate in the presence of TMAO

Differences were also seen in the amount of lactic acid produced by lactic acid bacteria in the presence and absence of TMAO (**Fig. 3B, D**). In raffinose-containing medium, bifidobacteria produced increased of amounts of lactate when grown in the presence of TMAO (the bifidobacteria grew poorly, if at all, in glucose-containing media). Unlike the bifidobacteria, the *Streptococcaceae* and *Enterococcaceae* grew well in the glucose-containing medium and produced over 25 mM lactic acid in the TMAO-containing samples compared with <5 mM in the control samples (**Fig. 3D**). To the best of our knowledge, this is the first time TMAO has been shown to influence the metabolism of gut bacteria – specifically lactic acid bacteria – without producing appreciable amounts of TMA.

## DISCUSSION

TMAO is a circulating metabolite produced as a direct result of microbial degradation of dietary methylamines in the intestinal tract, and can be readily detected along with its precursor TMA in human urine, blood and skeletal muscle [8,12,18,22]. It worsens atherosclerosis in some mouse models of CVD, and is positively correlated with CVD severity in humans. Beneficial effects associated with TMAO include potential protection from hyperammonia and glutamate neurotoxicity, alleviation of endoplasmic reticulum stress and improved glucose homeostasis by stimulating insulin secretion by pancreatic cells [10,15,16].

Over many decades it has been established that choline, PC and carnitine are dietary methylamines that contribute directly to microbiome-associated circulating levels of TMAO found in humans and other animals – e.g. [2,12,18,19,21,22]. However, TMAO itself is a water-soluble osmolyte found in high abundance in fish. Based on their observations that individuals with trimethylaminuria could reduce an oral dose of TMAO to TMA could not detoxify it to TMAO, al-Waiz *et al.* [25] suggested the gut microbiota could use TMAO as a substrate, and in those individuals without trimethylaminuria the TMA was re-oxidized in the liver before being excreted in the urine. This led these authors to propose the process of ‘*metabolic retroversion*’. Reference to the literature associated with fish spoilage and recent mining of metagenomic data have predicted members of the *Enterobacteriaceae* (particularly *Escherichia coli* and *Klebsiella pneumoniae*) have the potential to convert TMAO to TMA in the intestinal tract, but this has not been tested *in vitro* or *in vivo* to date [4,8,26–28]. Consequently, we instigated this study to demonstrate metabolic retroconversion of TMAO, and to determine the effect of TMAO on the growth and metabolism of human-derived intestinal bacteria in pure and mixed cultures.

Through *in vivo* administration of deuterated TMAO to mice via oral gavage in mice, we unambiguously demonstrated that TMAO is converted to TMA, with this TMA detectable in plasma within 2 h of administration. This conversion was highly dependent on the gut microbiota, as conversion of TMAO to TMA was dramatically reduced when the microbiota was suppressed by treatment of animals with broad-spectrum antibiotics. Even in the presence of antibiotics, there was low-level conversion of TMAO to TMA, suggesting a subpopulation of the microbiota was resistant to the antibiotics used in our experiment. However, administration of a broad-spectrum antibiotic cocktail for 14 days has been demonstrated to be an effective means of suppressing the gut microbiota in studies associated with gut microbial use of dietary methylamines [12].

Our *in vivo* experiment clearly demonstrates the gut microbiota converts TMAO to TMA, and that this TMA is re-oxidized to TMAO, in line with the process of ‘*metabolic retroversion*’ [25]. Gut-associated microbial conversion of the majority of TMAO to TMA is at odds with recent findings [8], in which it was suggested TMAO is taken up intact and not metabolised by the gut microbiota of humans. The ratio of TMAO-to-TMA is around 10:1 in humans and 1:10 in mice, meaning TMAO was *N*-oxidized before Taesuwan *et al.* [8] were able to observe it in circulating blood of their human subjects. In a human system with high FMO3 activity, ^17^O-labelled TMAO would need to be used in any study evaluating O turnover to allow calculation of the true rate of retroconversion.

Based on work done in mice [10,36], metabolic retroconversion of TMAO may be protective, and may even go some way to explaining why TMA and TMAO are detected at low levels in urine in the absence of dietary methylamines [17]. Chronic exposure of high-fat-fed mice to TMAO reduced diet-associated endoplasmic stress and adipogenesis, and improved glucose tolerance [10]. Exposure of high-fat-fed mice to TMA reduced low-grade inflammation and insulin resistance via inhibition of interleukin-1 receptor-associated kinase 4 (IRAK-4) [36]. Concentrations of the microbial signalling metabolite TMA used by Chilloux *et al.* [36] were comparable to those found in normal human plasma. Continual exposure of the human system to TMA/TMAO may bring similar benefits to host metabolic and inflammatory responses; these remain to be studied.

Having conducted our *in vivo* experiment in mice, we examined the ability of a range of human-derived gut bacteria to convert TMAO to TMA. In the mixed microbiota system and in pure cultures, the growth of the *Enterobacteriaceae* – the main TMA-producers – was quickly affected by the presence of TMAO. This is likely to happen in the human gut also. Consequently, we believe dietary TMAO undergoes metabolic retroconversion in mammals, with the TMA produced as a result of bacterial activity in the gut available to the host for conversion back to TMAO by FMOs in hepatocytes. It should be noted that [8] did not suppress or monitor the intestinal/faecal microbiota when they administered isotopically labeled TMAO to humans, nor did they measure d_9_-TMA and d_9_-TMAO in the portal vein, bypassing subsequent hepatic *N*-oxidation of d_9_-TMA, so it is not possible to interpret their results in the context of presence/absence of microbial activity.

Of note is the finding that caecal isolates of enterobacteria produce more TMA from TMAO than faecal isolates of the same bacterium. The speed with which TMAO was reduced to TMA by the *Enterobacteriaceae* in the present study suggests bacterial conversion of TMAO to TMA takes place in the small intestine/proximal colon of humans and small intestine/caecum of mice. It is, therefore, unsurprising that caecal bacteria – representing the microbiota present at the intersection of the small and large intestine – are metabolically more active than their faecal counterparts with respect to TMAO metabolism. This finding is relevant to functional studies of the gut microbiota where gene annotations are based largely on faecal isolates, whose functionalities may be greatly different from those of bacteria in other regions of the human intestinal tract [37,38]. It has already been demonstrated that the microbiota of the small intestine is enriched for functions associated with rapid uptake and fermentation of simple carbohydrates compared with the faecal microbiota, and that streptococci isolated from this niche are functionally very different from the same bacteria isolated from different habitats [37,38]. It is, therefore, important we characterize the functions and genomes of bacteria isolated from all regions of the intestinal tract, not just those of faecal bacteria, to gain a true picture of how microbial activity influences host health.

*Enterobacteriaceae* made the greatest contribution to the conversion of TMAO to TMA, both in pure culture and in a mixed microbiota. These Gram-negative bacteria are a source of the virulence factor lipopolysaccharide (LPS), which is associated with low-grade inflammation in high-fat-fed mice and elevated plasma levels that define metabolic endotoxaemia [39,40]. High-fat feeding has been shown to increase the representation of *Enterobacteriaceae* in the caecal microbiota of obesity-prone Sprague–Dawley rats [41], though this mode of feeding is known to modulate the microbiota of mice independent of obesity [42]. Non-LPS-associated virulence of *Enterobacteriaceae, Vibrio cholerae* and *Helicobacter pylori* is increased when these bacteria are grown anaerobically or microaerophilically in the presence of as little as 5–10 mM TMAO [43–46], and may be an additional means by which the gut microbiota contributes to CVD and other diseases in which increased representation of *Gammaproteobacteria* is observed. This warrants attention in future animal studies.

Lactic acid bacteria clearly grow better in the presence of TMAO. The relatively high concentration (1 %) of TMAO used in this study may have contributed to this improved growth, as TMAO is an osmolyte that stabilizes proteins. Future work will involve growing lactic acid bacteria in a range of TMAO concentrations to determine how this compound affects their growth and gene expression, and comparing faecal and caecal isolates. Similar to the *Enterobacteriaceae*, a large number of these bacteria are facultative anaerobes able to grow over a range of conditions, and whose representation is increased in obese, and cirrhotic patients [47,48]. *Streptococcus* and *Enterococcus* spp. are commensal lactic acid bacteria of the gut microbiota known to modulate immune function; but little is known about their metabolic activities in mixed microbial populations [49]. Understanding how commensal lactic acid bacteria influence the host in dysbiosis in mixed microbial communities may allow the development of approaches to modulate their activity and influence host health.

With respect to the lactic acid bacteria, it is important to note that our mixed culture work did not highlight these as being relevant to TMAO metabolism. This is unsurprising given these bacteria do not produce large quantities of TMA from TMAO. However, we have shown their metabolism is affected by presence of TMAO in growth medium, and the increased lactate they produce in its presence may contribute to cross-feeding associated with short-chain fatty acid production [50]. It is difficult to determine relevance of correlations from mixed microbial ecosystems in the absence of isotope labelling or pure culture work: i.e. correlation does not equate with causation. As an example, three (*Clostridium* clusters I and II, bifidobacteria, coriobacteriia) of the four groups of bacteria correlated with TMA production in our fermentation study did not produce notable quantities of TMA from TMAO based on our pure culture work. Therefore, correlating microbiota and metabolite data derived from complex systems will not give a true picture of which members of microbiota contribute to specific metabolic processes, and work with pure cultures is required to supplement functional studies to increase our understanding of poorly understood microbially driven metabolic processes within the human gut.

## CONCLUSIONS

We have demonstrated metabolic retroconversion – another example of host–microbial co-metabolism – occurs in the mammalian system with respect to TMAO, whereby TMAO is reduced by the gut microbiota to TMA and regenerated by host hepatic enzymes. We have also demonstrated that growth and metabolism of members of the gut microbiota are affected by TMAO in a source- and taxon-dependent manner, with the family *Enterobacteriaceae* making the greatest contribution to production of TMA in the gut.

## ABBREVIATIONS

AUC: area under the curve;
DMA: dimethylamine;
FISH: fluorescence *in situ* hybridization;
FMO: flavin mono-oxygenase;
MMA: monomethylamine;
PC: phosphatidylcholine;
TMA: trimethylamine;
TMAO: trimethylamine *N*-oxide;
UPLC–MS/MS: ultra-performance liquid chromatography–tandem mass spectrometry.

## DECLARATIONS

### Ethics approval

All animal procedures were authorized following review by the institutional ethics committee (Sorbonne Universities) and carried out under national license conditions. Ethical approval to collect caecal effluent from patients was obtained from St Thomas’ Hospital Research Ethics Committee (06/Q0702/74) covering Guy’s and St Thomas’ Hospitals, and transferred by agreement to London Bridge Hospital. Patients provided written consent to provide samples.

### Consent for publication

Not applicable.

### Availability of data and material

The datasets used and/or analysed during the current study are available from the corresponding authors upon reasonable request.

### Competing interests

The authors have no competing interests.

### Funding

MLJP was sponsored by Fundación Alfonso Martin Escudero (Spain). Work was supported by a grant from the EU-FP7 (METACARDIS, HEALTH-F4-2012-305312) to DG and MED. LH is in receipt of an MRC Intermediate Research Fellowship in Data Science (grant number MR/L01632X/1, UK Med-Bio).

### Authors’ contributions

MLJP, LH and ALM performed the microbiological work, JC was responsible for NMR analysis and interpretation of the NMR data, TA and FB did the animal work, MED supervised the NMR work, ALM supervised the microbiological work, CM provided invaluable advice for the labelling study and DG supervised the animal work. All authors contributed to the writing of the manuscript.

## Acknowledgements

Not applicable.

